# Gender, interdisciplinary graduate training, and confidence working in teams

**DOI:** 10.1101/2024.03.30.587416

**Authors:** Lesa Tran Lu, Laura Palucki Blake, Josh R. Eyler, Rafael Verduzco, Sibani Lisa Biswal, George N. Bennett, Jonathan J. Silberg

## Abstract

Teamwork is recognized as critical to solving complex societal challenges related to energy, health, and sustainability. With graduate education, students often gain teamwork experience through a problem-focused approach where they are brought into existing collaborations to pursue research that is focused on studying questions that have already been identified. Here, we describe an interdisciplinary educational program where graduate students were tasked with leading team formation, problem identification, and research formulation. This “team-first” training approach used a two-year curriculum to bring together students enrolled in diverse engineering and science graduate programs and provided students with a pedagogical understanding of interdisciplinarity, nurtured the development of student communication skills across disciplines, fostered student-led team formation and idea development, and empowered students to forge new connections between research groups. Assessment of three cohorts immediately following curriculum completion (n = 36) revealed significant gains in confidence in teamwork (p < 0.001) when compared to a control group of academic peers (n = 74). These gains varied across demographic groups, with women in science, technology, engineering, and mathematics presenting the strongest gains. This finding illustrates the importance of exploring how interdisciplinary team curricula in graduate school could support overcoming the gender gap in confidence.

**Significance:** Pedagogical models for graduate education often neglect the importance of teamwork training. Here, we describe an interdisciplinary training program that was developed to bring together doctoral students from diverse science, technology, engineering, and mathematics disciplines for a two-year curriculum that focused on teamwork training through student-led team formation, problem identification, and research formulation. Following program participation, we measured participant confidence in teamwork relative to a peer group. Our findings reveal gains with confidence in teamwork, with women presenting the strongest increases without negative effects on other groups. This pedagogical approach represents a strategy to close gender gaps in professional role confidence while complementing the benefits of traditional disciplinary training approaches.

## Introduction

Society is facing challenges of growing complexity related to energy (1), sustainability (2), and health (3), whose solutions necessitate collaboration across disciplinary and organizational structures (4). In parallel, technological breakthroughs in data science (5), materials science (6), and synthetic biology (7) are creating new opportunities for collaborations across disciplines to transform industries as we work to discover solutions to these global challenges. To fertilize such connections, organizations are increasingly turning to team-based structures as a mechanism to innovate and adapt to complex and dynamically changing societal-technical needs (8). This shift in organizational structure has led to the recognition of the importance of studies that examine team effectiveness models and an appreciation of the importance of teamwork training (9, 10).

Employers identify teamwork as a key skill when searching for job candidates. In a recent survey of more than one thousand employers, over 75% reported that they put value on job candidates with skills to work effectively in teams, to work with people from different backgrounds, and to take initiative (11). Additional skills identified as very important included strong oral and written communication, creative and innovative thinking, and an ability to integrate and apply ideas across different settings and contexts (11). Over half of those surveyed felt that effective workers require knowledge gained from thinking across disciplines, *i.e.*, interdisciplinary thinking, and more than two-thirds noted that success in the workplace benefits from a mindset that is comfortable engaging with people who have a diverse range of backgrounds, opinions, and viewpoints (11). Similarly, a job outlook study conducted by the National Association of Colleges and Employers, which surveyed over three hundred companies, revealed that a similar percentage of employers seek candidates with strong teamwork skills (12). These two reports highlight the importance of connecting workplace readiness, defined by employer needs, and academic preparation as the costs of undergraduate and graduate education continue to grow (13, 14).

As national agencies and organizations have emphasized the value of teamwork (15, 16), universities have focused on developing pedagogical models that train students in interdisciplinary research and teamwork (17). In line with the high-impact teaching practices outlined by the American Association of Colleges and Universities (18), these efforts have invested in expanding curricular opportunities for collaborative projects through the lens of a “big theme or question” that emphasizes the importance of teamwork in addressing complex real-world challenges (19–21). At the graduate level, the National Science Foundation (NSF) has provided support to develop pedagogical approaches for interdisciplinary training via the Integrative Graduate Education and Research Traineeship (IGERT) and NSF Research Traineeship (NRT) programs, which seek to effectively train graduate students in convergent research areas to better align with the needs of a dynamically changing workforce (22). Innovative models for interdisciplinary graduate training have been developed through these programs, with many sharing an emphasis on exploration, communication, and collaboration across disciplines (23–27).

With most IGERT and NRT training programs, a problem-focused training approach is implemented where students join teams that are unified around previously identified research challenges (28, 29). However, findings suggest that students should concurrently develop their interdisciplinary research methods and collaborative meta-competencies, including their networking, epistemic adaptability, and value assessment skills (30–33). One recent approach targeted these skills by tasking students with leading team formation, problem identification, and research formulation (34). While this “team-first” pedagogical model achieved integration among students, there was limited assessment of skills crucial for workforce preparedness as perceived by employers (35). Here, we describe a team-first interdisciplinary graduate training program centered on scholarship at the cell-material interface and an assessment of student creativity and confidence in teamwork following program participation.

## Results and Discussion

### The interdisciplinarity curriculum

The curriculum, which spanned four semesters, was open to students from diverse engineering, natural sciences, and social sciences doctoral programs. This program was designed to provide students with a pedagogical understanding of teamwork, practical experience communicating across disciplines, and open-ended experiences with forging new research connections. With distinct pedagogical goals for each semester, this curriculum sought to foster student creativity and confidence in teamwork by encouraging regular student brainstorming (36), allowing students to experiment with ideas that cross disciplinary boundaries (37), generating opportunities for passion projects (38), embracing a diversity of skill sets in teams (39), providing a safe space for creative endeavors (40), and rewarding innovation through seed funds for student-led projects (41). To minimize curriculum burdens, no more than two credit hours of coursework was required each semester. An overview of the curriculum is summarized in Figure 1.

**Figure 1.**
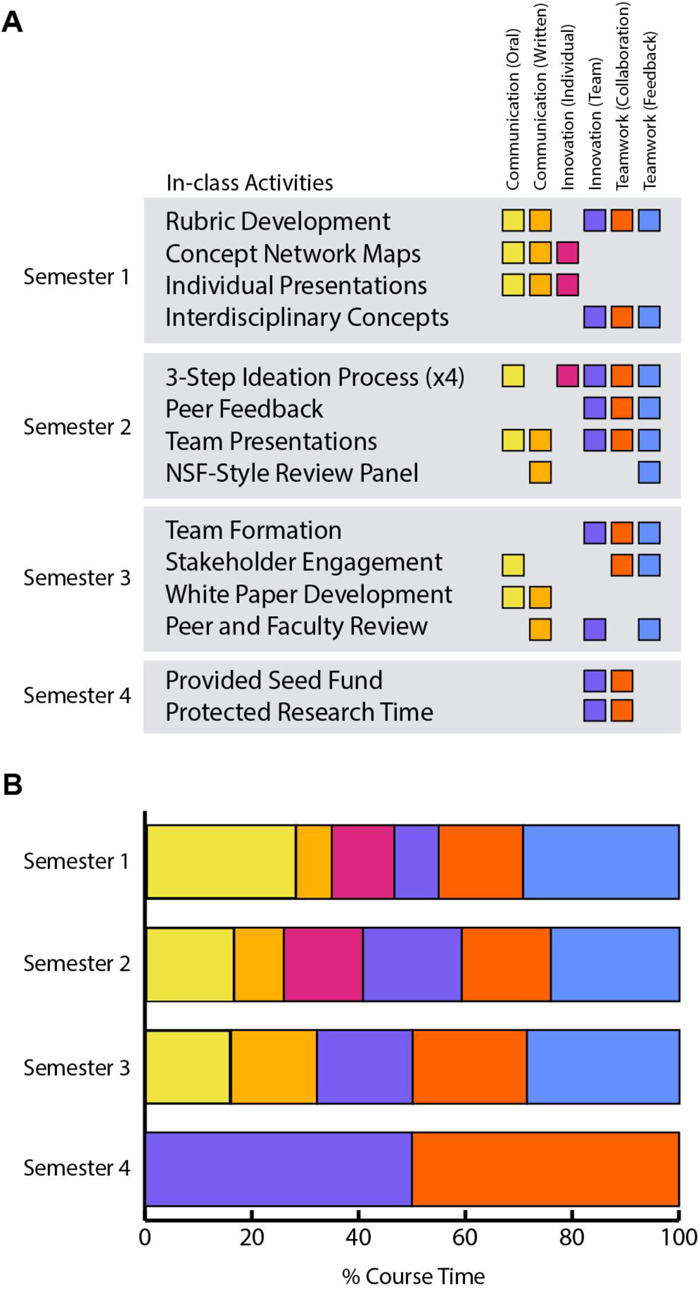
Curriculum built for teamwork training. (**A**) The activities conducted during each semester to support graduate student skill building in communication, innovation, and teamwork. Each semester had a major pedagogical emphasis, with the first focused on building familiarity, the second focused on fostering creativity, the third focused on developing team research projects, and the fourth focused on pursuing collaborative research. (**B**) The four-semester curriculum varied in the time devoted to developing communication, teamwork, and innovation skills. To minimize the overhead of the curriculum, activities were largely conducted during class time.

**Figure 2.**
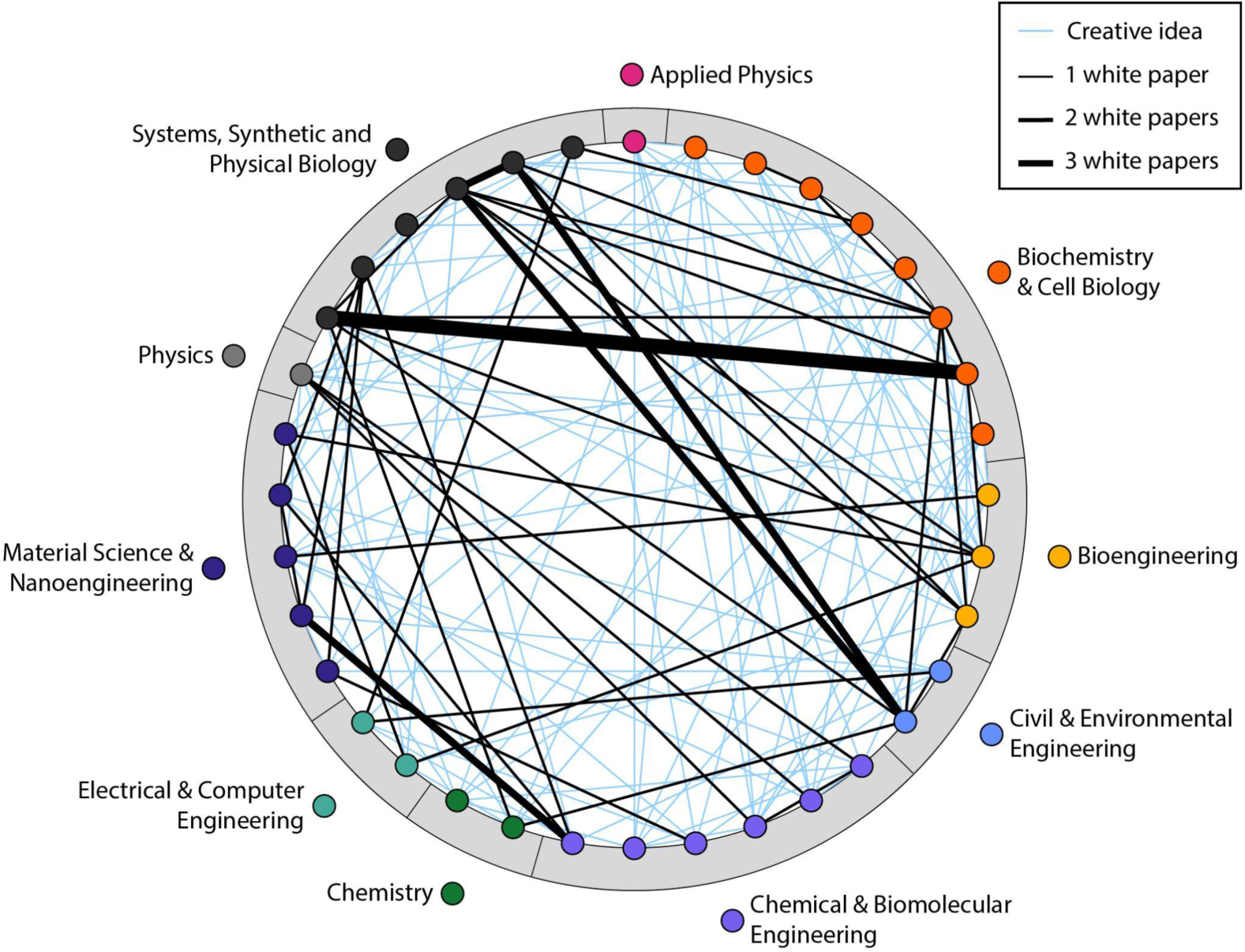
Network map illustrating student connections. The network illustrates the diversity of research ideas generated by student teams across the three program cohorts, which included 36 students training across ten doctoral programs. Nodes represent students, color-coded by training program. Creative ideas developed as part of the second-semester curriculum are shown in blue (n = 47). With this curriculum, students were charged with generating interdisciplinary ideas that required the expertise of two or more trainees. The black lines represent teams formed by students as part of the third-semester curriculum when they developed white papers around ideas that they wanted to pursue (n = 19). The only constraint on these white papers was that students make a new connection between two or more labs in a way that did not exist previously. Line thickness represents the number of white papers collaboratively developed.

The first-semester curriculum was designed to help students from different disciplines build familiarity with one another, learn how to respond to questions from individuals outside of their discipline, and gain confidence in communication in this interdisciplinary setting. To achieve these goals, students were asked to collaboratively develop a rubric for evaluating effective communication across disciplines. Students then used this rubric to guide the development and completion of student-led activities, including a tour of a piece of research equipment in each student’s laboratory and short presentations that introduced their research. With the latter activity, the time protected for questions and discussion was equivalent to the time allotted for presentations, similar to a Gordon Research Conference presentation (42). To gain a pedagogical understanding of team concepts, students discussed different interdisciplinarity concepts from assigned reading (43).

The second-semester curriculum was designed to foster student creativity and build student confidence in pitching new research ideas. Our operational definition of creativity when developing this curriculum was the ability to generate an innovative research idea that could have value and is non-obvious to a scholar trained in a single discipline (44, 45). To achieve these goals, students participated in a three-step ideation process, which was repeated four times in total. First, each student presented a review article of relevance to the assigned topic (*e.g.*, bioelectronics and health), a peer-reviewed research article that inspired their idea, and a brief description of their idea. Then, students were assigned to small groups with members from two or more doctoral programs. These groups were tasked with discussing their individual ideas, identifying a single idea to further develop as a team, and developing a research plan that allowed each team member to contribute to the innovation. Finally, each group presented their ideas to the class, who were tasked with asking questions, providing additional feedback, and logging their thoughts on each idea in a shared document. At the end of the semester, students evaluated a subset of the ideas generated over the semester via a mock review panel that used the NSF merit review criteria.

The third-semester curriculum was designed to foster student creativity, to grow confidence in leading team formation, and to grow confidence in writing proposals. To achieve this, students independently developed one-page idea plans which included their research idea and proposal for team composition and synergies. Students then shared these documents with their peers to gather feedback on potential collaborator interest. To ensure interdisciplinary team formation, all teams were required to have members with different thesis advisors and to have not previously collaborated on a research project. The students then worked in teams to develop an interdisciplinary research proposal in the format of a white paper, which provided background information, an experimental plan, and a description of team synergies. These white papers were reviewed by a faculty committee, who each provided feedback on the hook, team, plan, benefits, and investment. Once approved, teams were awarded seed grants totaling up to $3,000 to pursue the idea.

The final semester of the curriculum provided student teams with protected time to pursue their collaborative research idea. This activity was designed to help students develop confidence through practical teamwork by allowing them to pursue the experimental plans outlined in their white papers.

### Offering the curriculum

Similarly sized student cohorts with 11 to 14 students began the curriculum in 2019, 2020, and 2021. Each cohort included students enrolled in at least six different graduate programs, with a total of ten training programs being represented across all three cohorts. Students were pursuing doctoral degrees in disciplinary programs within five departments across the schools of engineering (Bioengineering; Civil and Environmental Engineering; Chemical and Biomolecular Engineering; Electrical and Computer Engineering; Materials Science and Nanoengineering) and three programs within natural sciences (Biochemistry and Cell Biology; Chemistry; Physics). Additionally, students were recruited from a pair of interdisciplinary programs (Applied Physics; Systems, Synthetic and Physical Biology) that connect departments within these two schools. A subset of students were offered a stipend to participate in the program. In the 2019 cohort, six students received two years of funding, while the other five were non-funded participants. Non-funded students represented a mix of domestic and international students; the latter were not eligible for funding based on NSF rules. Two of the non-funded participants received stipends during their second year in the program. For the 2020 and 2021 cohorts, funding was conditional upon successful completion of the first-year curriculum, thereby allowing a greater number of students to be funded initially. In total, 78% of the participants across the three cohorts received funding. Only one student withdrew from the program, which occurred after they completed the first year of the curriculum. Upon completion of the program, each student was asked to identify 3 non-participating peers in their respective graduate programs who would serve as part of the control group. Table 1 provides the demographics for students completing the program, the demographics for the control group used in the assessment, and the graduate programs where students were enrolled.

**Table 1.** Student cohort information and survey respondent demographics. In total, thirty-six trainees completed the program and the survey (100%). Each trainee provided a list of three peers outside the program, who were also asked to complete the survey. Of the 108 peers who served as controls, 74 completed the survey (69%).

|  |  | CONTROL |  | TRAINEES |  |
| --- | --- | --- | --- | --- | --- |
| School/Program |  | Number | Percent | Number | Percent |
| School of Engineering (ENGI) |  |  |  |  |  |
|  | Bioengineering | 9 | 12% | 4 | 11% |
|  | Civil and Environmental Engineering | 5 | 7% | 3 | 8% |
|  | Chemical and Biomolecular Engineering | 8 | 11% | 5 | 14% |
|  | Electrical and Computer Engineering | 6 | 8% | 2 | 6% |
|  | Materials Science and Nanoengineering | 9 | 12% | 5 | 14% |
| School of Natural Sciences (NSCI) |  |  |  |  |  |
|  | Biochemistry and Cell Biology | 15 | 20% | 7 | 19% |
|  | Chemistry | 5 | 7% | 2 | 6% |
|  | Physics | 2 | 3% | 1 | 3% |
| Interdisciplinary Programs (ENGI and NSCI) |  |  |  |  |  |
|  | Applied Physics | 2 | 3% | 1 | 3% |
|  | Systems, Synthetic and Physical Biology | 13 | 18% | 6 | 17% |
| **Demographics** |  |  |  |  |  |
| Gender |  |  |  |  |  |
|  | Men | 34 | 46% | 18 | 50% |
|  | Women | 40 | 54% | 17 | 49% |
|  | missing |  |  | 1 | 1% |
| Race |  |  |  |  |  |
|  | Asian | 24 | 32% | 8 | 22% |
|  | Black or African American | 8 | 11% | 1 | 3% |
|  | Hispanic or Latinx | 11 | 15% | 2 | 6% |
|  | Middle Eastern/N. African | 3 | 4% | 0 | 0% |
|  | White | 24 | 32% | 17 | 47% |
|  | Two or more races | 4 | 5% | 7 | 19% |
|  | Unknown | 0 | 0% | 1 | 3% |
| Historically Underrepresented Group |  |  |  |  |  |
|  | Yes | 48 | 65% | 20 | 56% |
|  | No | 26 | 35% | 16 | 44% |
| First Generation |  |  |  |  |  |
|  | Yes | 8 | 11% | 5 | 14% |
|  | No | 65 | 89% | 31 | 86% |
| International Student |  |  |  |  |  |
|  | Yes | 38 | 51% | 5 | 14% |
|  | No | 36 | 49% | 31 | 86% |

The coursework requirements were identical for each cohort, with a similar profile of products generated by each group. In the first semester, each student developed and presented a concept network map on their research interests. Students also gave short oral presentations on their thesis research. Students also collaborated to create a rubric for assessing communication across disciplines and to generate a single network map to orient student interests and potential research connections. In the second semester, each student developed multiple research ideas independently. In addition, they were assigned to small interdisciplinary teams to collaboratively develop ideas. In the third semester, students drafted white papers proposing a new interdisciplinary research collaboration. These white papers, which represented the capstone team synthesis activity in the curriculum, connected students across two or more doctoral programs. In the final semester, students pursued their proposed collaboration, with a modest amount of financial support from the program.

The first cohort reported that the idea development in the second semester was stressful. This led to the addition of four activities to the first-semester curriculum to familiarize students with team-based ideation. First, a low-stakes idea generation activity was added at the beginning of the semester, during which more senior program participants collaborated with new students to pitch ideas. Next, new students were invited to attend presentations given by more senior students who were showcasing their white papers in their third semester. These presentations were designed to mimic the Faraday Discussions coordinated by the Royal Society of Chemistry (46), where individuals attending the presentations have reviewed the white papers and record their feedback in advance, leaving ample time to discuss following the presentation. Immediately following the white paper presentations, the senior students participated in a panel discussion that was designed to educate the new students about the white paper development process. Lastly, a low-stakes idea generation activity was implemented at the end of the first semester to provide new students a chance to rapidly ideate in a single class session prior to the second semester. While the curriculum was designed to utilize in-class interactions, some of the coursework was delivered virtually during the COVID-19 pandemic in the spring and fall 2020.

Several synergistic activities were organized beyond the curriculum. To help students develop a broad perspective about the university beyond their training programs, we organized a boot camp prior to the coursework, which involved round table discussions with the Technology Transfer Office, the Doerr Institute for New Leaders, and the Baker Institute for Public Policy. To provide students with leadership experience, they were charged with working in teams to create committees with a topical focus, such as seminars, recruiting, and mental wellbeing committees. Students generated proposals for annual activities with a mission statement and a budget, pursued the proposed activities with financial support from the program, and reported on their efforts annually.

### Assessing confidence and creativity

Our team-first program was designed to help students learn how to collaborate across disciplines. We hypothesized that this curriculum would increase student confidence in teamwork, foster creativity, and support confidence in graduate school. These ideas were evaluated by asking trainees to complete a survey following program completion. The survey was also administered to non-participating graduate student peers who were identified by trainees.

Looking at all three cohorts immediately following program completion, trainees (24.69) showed significantly more confidence in teams (t = 3.22, p<0.001) compared to their peers in the control condition (28.89). A closer look at this finding reveals that the difference is most prominent among women (trainees = 25.24, controls = 31.90, t = 3.85, p <0.001). Table 2 summarizes the significant results. Among the items that make up this assessment tool, eight of the twelve items with significant differences (75%) were in the hypothesized direction. Trainees were more likely than their peers in the control condition to agree with the following statements: (i) I am comfortable initiating conversations with researchers outside my primary discipline, (ii) I am quite good at talking about my research ideas in ways that bring other researchers into the conversation, (iii) I am able to identify potential contributors who can move a research idea forward than controls, and (iv) I am comfortable leading a group of my academic peers than controls. In addition, trainees were more likely than their peers in the control condition to disagree with the following statements: (v) I follow my advisor’s lead and let them do the networking than controls, (vi) sometimes, I’m not sure how to contribute to (inter) disciplinary conversations, (vii) I do my best research when they work alone, and (viii) I hesitate to be involved in projects that require me to work on a team where I am the only member in my discipline. A closer look at confidence in teams shows that for seven of the eight significant findings (87.5%), trainees who were members of groups historically underrepresented in STEM, which included women, were significantly higher than their control peers. This observation suggests that while confidence in teams is higher for trainees overall than for controls, trainees who are members of groups historically underrepresented in STEM disproportionally benefit when compared to their peers who did not participate in the interdisciplinary curriculum.

**Table 2.** Significant trends for confidence in teamwork. The average scores for all twelve questions are provided as well as the eight questions where significant differences were observed between the participant and the control groups. Difference values are calculated with more significant digits than what is shown, and statistics are from two-sample t tests. HUG = historically underrepresented groups in STEM. Items that were reverse coded are noted with an (R).

|  | All Respondents |  | Men |  | Women |  | HUG |  | First Generation |  |
| --- | --- | --- | --- | --- | --- | --- | --- | --- | --- | --- |
|  | Control (n = 74) |  | Control (n = 34) |  | Control (n = 40) |  | Control (n = 48) |  | Control (n = 8) |  |
|  | Trainee (n = 36) |  | Trainee (n = 18) |  | Trainee (n = 17) |  | Trainee (n = 20) |  | Trainee (n = 5) |  |
|  | Means | Sig | Means | Sig | Means | Sig | Means | Sig | Means | Sig |
| Confidence in Teamwork Scale | C = 28.89 | p<.001 |  | n/a | C = 31.90 | p<.001 | C = 30.77 | p<.001 |  | n/a |
|  | T = 24.69 |  |  |  | T = 25.24 |  | T = 25.60 |  |  |  |
| I am comfortable initiating conversations with researchers outside of my primary discipline | C = 2.11 | p<.05 | C = 1.97 | p<.05 |  | n/a | C = 2.23 | p=.062 |  | n/a |
|  | T = 1.72 |  | T = 1.56 |  |  |  | T = 1.85 |  |  |  |
| Sometimes, I'm not sure how to contribute to (inter) disciplinary conversations (R) | C = 2.61 | p<.05 |  | n/a | C = 2.20 | p>.01 | C = 2.38 | p<.05 |  | n/a |
|  | T = 3.08 |  |  |  | T = 3.00 |  | T = 2.95 |  |  |  |
| I'm quite good at talking about my research ideas in ways that bring other researchers into the conversation | C = 2.61 | p<.05 |  | n/a | C = 2.83 | p<.01 | C = 2.69 | p<.05 |  | n/a |
|  | T = 2.25 |  |  |  | T = 2.06 |  | T = 2.20 |  |  |  |
| I would rather follow my advisor's lead and let them do the networking (R) | C = 3.09 | p<.01 | C = 3.53 | p=.05 | C = 2.73 | p<.01 | C = 2.79 | p<.01 |  | n/a |
|  | T = 3.78 |  | T = 4.00 |  | T = 3.53 |  | T = 3.55 |  |  |  |
| I can identify potential contributors who can move a research idea forward | C = 2.28 | p<.05 |  | n/a | C = 2.50 | p<.01 | C = 2.42 | p<.05 |  | n/a |
|  | T = 1.89 |  |  |  | T = 1.76 |  | T = 1.85 |  |  |  |
| I hesitate to be involved in projects that require me to work on a team where I am the only member in my discipline (R) | C = 3.70 | p<.01 |  | n/a | C = 3.50 | p<.001 |  |  |  | n/a |
|  | T = 4.25 |  |  |  | T = 4.41 |  |  |  |  |  |
| I do my best research when I work alone (R) | C = 3.10 | p<.01 |  | n/a | C = 2.90 | p<.01 | C = 3.00 | p<.01 | C = 2.63 | p<.05 |
|  | T = 3.67 |  |  |  | T = 3.94 |  | T = 3.90 |  | T = 4.00 |  |
| I am comfortable leading a group of my academic peers | C = 2.34 | p<.05 |  | n/a | C = 2.65 | p<.05 | C = 3.80 | p=.062 |  | n/a |
|  | T = 2.00 |  |  |  | T = 2.06 |  | T = 2.50 |  |  |  |

No significant differences were observed between trainee creativity and their peers in the control condition when considering the overall scale. However, a closer inspection of the individual items that make up the scale reveals that four of the fourteen items (28%) presented significant differences between the two groups (Table 3), all of which were in the predicted direction. Trainees who participated in the interdisciplinary curriculum were more likely than their peers in the control group to agree with the following statements: (i) I brainstorm new ideas beyond the ones my advisor gives me, (ii) I take the time to brainstorm new ideas beyond the ones my advisor gives me, (iii) I have a lot of intellectual curiosity, and (iv) I look for new research ideas outside my primary discipline. With all four items, women trainees were significantly higher than their control peers, while only three items were significantly higher with trainees from historically underrepresented groups (HUG) in STEM. Taken together, this data reinforces the idea that women and historically underrepresented groups make gains in creativity compared to their peers in the control condition.

**Table 3.**
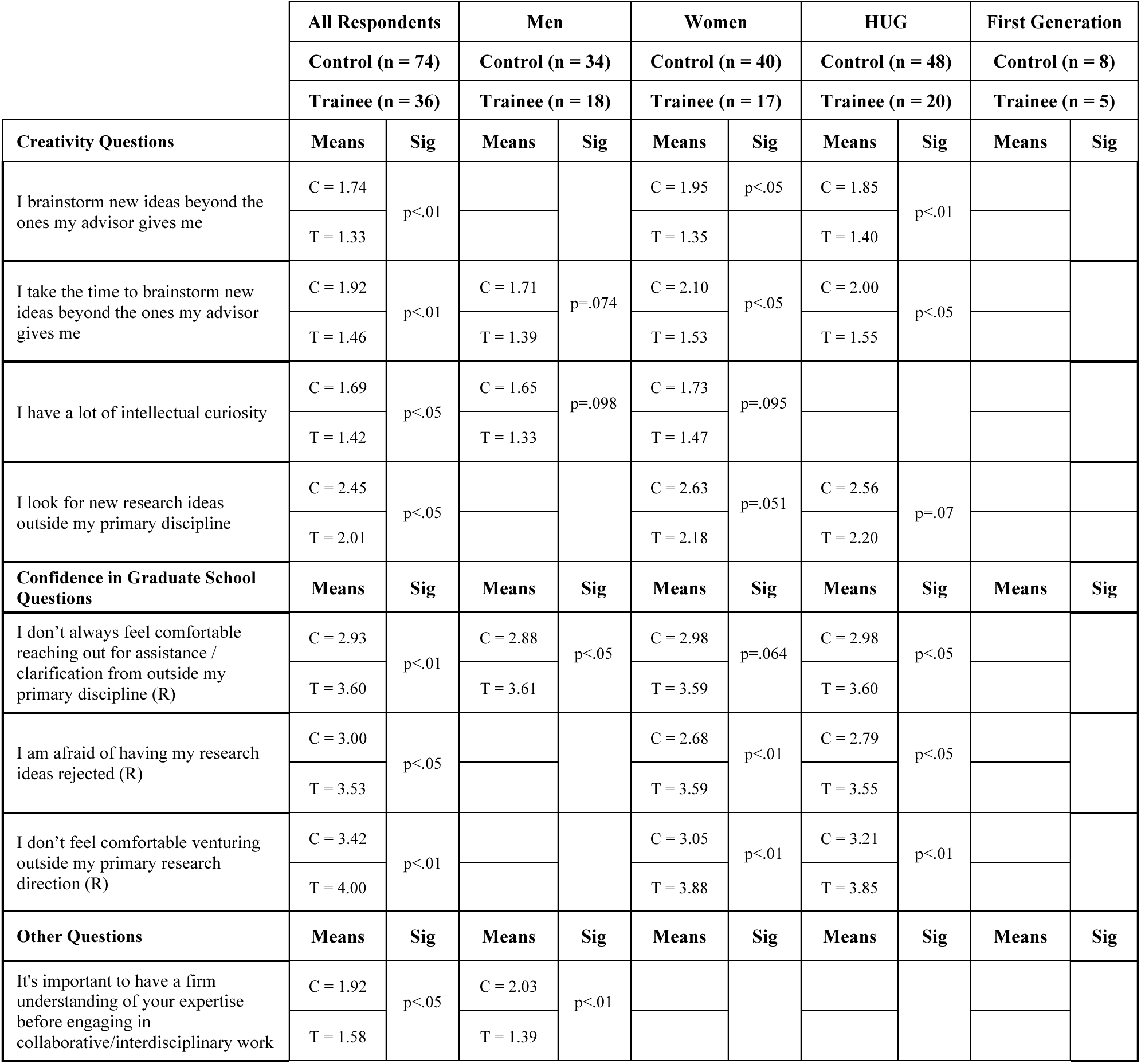
Significant trends for creativity and confidence in graduate school. The average scores for all questions were not significant but a subset of the questions in each tool presented significant differences between the participant and the control groups. Difference values are calculated with more significant digits than what is shown, and statistics are from two-sample t tests. HUG = historically underrepresented groups in STEM. Items that were reverse coded are noted with an (R).

|  | All Respondents |  | Men |  | Women |  | HUG |  | First Generation |  |
| --- | --- | --- | --- | --- | --- | --- | --- | --- | --- | --- |
|  | Control (n = 74) |  | Control (n = 34) |  | Control (n = 40) |  | Control (n = 48) |  | Control (n = 8) |  |
|  | Trainee (n = 36) |  | Trainee (n = 18) |  | Trainee (n = 17) |  | Trainee (n = 20) |  | Trainee (n = 5) |  |
| Creativity Questions | Means | Sig | Means | Sig | Means | Sig | Means | Sig | Means | Sig |
| I brainstorm new ideas beyond the ones my advisor gives me | C = 1.74 | p<.01 |  |  | C = 1.95 | p<.05 | C = 1.85 | p<.01 |  |  |
|  | T = 1.33 |  |  |  | T = 1.35 |  | T = 1.40 |  |  |  |
| I take the time to brainstorm new ideas beyond the ones my advisor gives me | C = 1.92 | p<.01 | C = 1.71 | p=.074 | C = 2.10 | p<.05 | C = 2.00 | p<.05 |  |  |
|  | T = 1.46 |  | T = 1.39 |  | T = 1.53 |  | T = 1.55 |  |  |  |
| I have a lot of intellectual curiosity | C = 1.69 | p<.05 | C = 1.65 | p=.098 | C = 1.73 | p=.095 |  |  |  |  |
|  | T = 1.42 |  | T = 1.33 |  | T = 1.47 |  |  |  |  |  |
| I look for new research ideas outside my primary discipline | C = 2.45 | p<.05 |  |  | C = 2.63 | p=.051 | C = 2.56 | p=.07 |  |  |
|  | T = 2.01 |  |  |  | T = 2.18 |  | T = 2.20 |  |  |  |
| Confidence in Graduate School Questions | Means | Sig | Means | Sig | Means | Sig | Means | Sig | Means | Sig |
| I don't always feel comfortable reaching out for assistance / clarification from outside my primary discipline (R) | C = 2.93 | p<.01 | C = 2.88 | p<.05 | C = 2.98 | p=.064 | C = 2.98 | p<.05 |  |  |
|  | T = 3.60 |  | T = 3.61 |  | T = 3.59 |  | T = 3.60 |  |  |  |
| I am afraid of having my research ideas rejected (R) | C = 3.00 | p<.05 |  |  | C = 2.68 | p<.01 | C = 2.79 | p<.05 |  |  |
|  | T = 3.53 |  |  |  | T = 3.59 |  | T = 3.55 |  |  |  |
| I don't feel comfortable venturing outside my primary research direction (R) | C = 3.42 | p<.01 |  |  | C = 3.05 | p<.01 | C = 3.21 | p<.01 |  |  |
|  | T = 4.00 |  |  |  | T = 3.88 |  | T = 3.85 |  |  |  |
| Other Questions | Means | Sig | Means | Sig | Means | Sig | Means | Sig | Means | Sig |
| It's important to have a firm understanding of your expertise before engaging in collaborative/interdisciplinary work | C = 1.92 | p<.05 | C = 2.03 | p<.01 |  |  |  |  |  |  |
|  | T = 1.58 |  | T = 1.39 |  |  |  |  |  |  |  |

Differences were also not observed between trainees and the control group on the confidence in graduate school scale. However, three of the eighteen (17%) items were significant (Table 3), all in the predicted direction. Following completion, trainees were more likely than their peers in the control condition to disagree with the following: (i) I don’t always feel comfortable reaching out for assistance/clarification from outside my primary discipline, (ii) I am afraid of having my research ideas rejected, and (iii) I don’t feel comfortable venturing outside my primary research direction. As was the case with confidence in teams and creativity items, women trainees and members of historically underrepresented groups in STEM were significantly higher than their peers in control conditions on all four items. In contrast, male trainees were significantly different from their peers in the control condition on only one item, *i.e.*, I don’t always feel comfortable reaching out for assistance/clarification from outside my primary discipline. The fact that for items across confidence in teams, creativity, and confidence in graduate school women and historically underrepresented groups in STEM make gains compared to their peers in the control condition again underscores the relevance of interdisciplinary training curriculum and pedagogy in graduate education in STEM.

### Implications

The team-first curriculum studied herein provided doctoral students an opportunity to augment their disciplinary training by regularly engaging with peers from different backgrounds, synthesizing a range of team ideas in an intellectually safe space, identifying an interdisciplinary innovation that they wanted to explore as part of a team, and pursuing that research idea with the support of seed funds from the program. By having students practice multiple skills that employers search for in job candidates (11, 12), ranging from oral and written communication to working effectively in teams and with people from different backgrounds to creative and innovative thinking, this curriculum led to significant gains in student confidence in teamwork. This outcome is well-aligned with one mindset that employers view as important for success in the workplace, namely comfort in engaging with people who have a diverse range of opinions and viewpoints (11). Surprisingly, women presented the strongest gain in confidence following participation in this curriculum. This observation suggests that the team-first curriculum may represent a strategy to overcome gender gaps in professional role confidence (47, 48), which has been posited to underlie the attrition of women in engineering career paths as well as the gender pay gap among science, technology, engineering, and mathematics graduates (49).

The underlying cause for the confidence gain observed with women following participation in the team-first training program is not known. One possible explanation is that the Dunning-Kruger effect underlies these trends (50, 51). The male students assessed may have been less aware of their deficiencies in teamwork. In the future, it will be interesting to explore which aspects of the curriculum are most important in contributing to the observed benefits, whether similar team-based education approaches can address gender gaps in confidence without negatively affecting other groups, and if the trends observed herein extend to female graduate students from different countries and other historically underrepresented groups in STEM that also present gaps in confidence (52). In addition, to better align with employer needs and support the development of self-confidence, which has been identified as a key leadership trait (53), it will be important to explore how traditional models of graduate student pedagogy can be re-oriented, refined, and scaled through the addition of cost-effective team-first activities to graduate coursework, cohort-based co-curricular research, or research labs.

## Materials and Methods

### Student recruitment

Three student cohorts were recruited to this curriculum between 2019 and 2021 by advertising this opportunity to students enrolled in a dozen different doctoral programs and conducting annual information sessions. The application asked students to provide basic information, such as their stage of training, thesis advisor status, coursework progress, and demographics. Also, short essays (<500 words) were required that described: (i) their interest in the interdisciplinary topic of bioelectronics, (ii) any relevant team experiences that influenced their interest, and (iii) how the training program could enhance their careers. All applicants were interviewed by a faculty committee that asked questions related to the essay topics and offered applicants a chance to ask questions. Using this information, the admissions committee admitted cohorts to achieve representation from at least six training programs. Offer letters were then provided to students which outlined the requirements of the program, including a commitment to participating in assessment. Students were only allowed to participate if they and their thesis mentor returned a signed offer letter acknowledging that they understood the full program requirements.

### Survey data collection

We employed a confidential survey, including both qualitative and quantitative elements, to determine to what extent program goals were being achieved. Trainees were asked to complete the survey upon completing the program, and to name at least three peers who were not in the program who could be invited to participate as controls. This strategy allows us to compare those in the program with those who are not at a single point in time. Participants who completed the survey were compensated with a $25 gift card. Response rates and survey participant demographics are provided in Table 1.

### Survey structure

For the construction of the scales, in addition to developing original items, we adapted items from Bauer’s Chemistry Self-Confidence Inventory (54), the Rosenberg Self-Esteem scale (55), and the scholarly creativity items from Kaufmann’s Domains of Creativity Scale (56). Bauer’s chemistry self-concept inventory consists of 5 subscales: mathematics self-concept, chemistry self-concept, academic self-concept, academic satisfaction self-concept, and creativity self-concept. We adapted nine items. Rosenberg’s self-esteem scale consists of ten items, the purpose of which is to measure self-esteem. Originally it was designed to measure the self-esteem of high school students. However, since its development, the scale has been used with a variety of groups including adults pursuing university studies (57), with norms available for many different groups. We adapted four items to measure confidence in graduate school. Kaufmann’s domains of creativity scale have five domains: Self/Everyday, Scholarly, Performance, Mechanical/Scientific, and Artistic. We adapted two items from the scholarly domain.

Through the adaptations made from these three instruments, plus the addition of our own items, we built three scales: Creativity (thirteen items; Table S1), Confidence in Teamwork (twelve items; Table S2), and Confidence in Graduate School (eighteen items; Table S3). Our choice to adapt items from each of the pre-existing scales was based on the fact that our population consisted of STEM graduate students broadly. We wanted the items to be modern and relevant to all participants, regardless of discipline. All Items employed a 5-point Likert-style scale with the following response options: Strongly Agree, Agree, Neutral, Disagree, and Strongly Disagree. For data processing purposes, we assigned values from 1 to 5, respectively. In our scales, there were inverted statements. In scoring these items, we reverse-coded them so that 1 = Strongly Disagree. While we observed the internal validity of our scales for our purposes, the authors stress the need for continued validation of each of these scales with different populations. In addition to the items in the three scales, the survey included open-ended responses to the following questions: (i) please describe the most significant learning experience you have had so far this year; (ii) what kinds of things (*e.g.*, mentoring, advising, switching lab, taking a leave) have impacted your confidence this year; (iii) what kinds of things have impacted your creativity this year; and (iv) if you had a magic wand, what additional graduate training opportunities to nurture your creativity and confidence would you create. The last item on the survey asked participants to share any additional comments regarding their educational experience this year. Finally, the survey collected the following demographic information: (i) department and training program, (ii) semester and year they started in their program, (iii) gender, (iv) race, (v) first generation, and (vi) international student status.

## Acknowledgments

We are grateful for support from the National Science Foundation under grant 1828869 and the Institute for Biosciences and Bioengineering and Rice University for administrative support of this program.

