## Supplemental Tables for "Gender, interdisciplinary graduate training, and confidence working in teams"

**Supplemental Table 1. Creativity survey.** In total, fourteen items were used to create this scale. Items that are reverse (R) coded are noted as well as the source of items. Cronbach's Alpha yielded a value of 0.728.

| Creativity Scale | Source |
|---|---|
| I brainstorm new ideas beyond the ones my advisor gives me. | |
| I take the time to brainstorm new ideas beyond the ones my advisor gives me. | |
| I have a lot of intellectual curiosity. | Bauer-original |
| I am good at combining research ideas in ways that others have not tried. | Bauer-adapted |
| I look for new research ideas outside my primary discipline and try to apply them to my own. | |
| I have gone back to research ideas I have rejected before | |
| In science, it is helpful to have people approaching a problem from different angles | |
| Sometimes the complexity of a research problem in my discipline demands diversity of thought. | |
| When I have a new research idea, I discuss it with others to evaluate its potential. | |
| When I am generating new research ideas, I try not to evaluate them until I have generated many ideas. | |
| I am able to prioritize which research ideas to pursue. | |
| I tend to lose my sense of time while I'm engaged in my research. | |
| Carrying out or designing a scientific experiment. | Kauffman-adapted |
| Attempting to explain an idea using different methods (*e.g.*, visually, anecdotally) so that different people can understand it. | Kauffman-adapted |

**Supplemental Table 2. Confidence in teamwork survey.** In total, twelve items were used to create this scale. Items that are reverse (R) coded are noted as well as the source of items. Cronbach's Alpha yielded a value of 0.783.

| Confidence in Teamwork Scale | Source |
|---|---|
| I am comfortable asking questions in order to understand my peer's research ideas. | |
| I am comfortable initiating conversations with researchers outside of my primary discipline. | Bauer-adapted |
| Sometimes, I'm not sure how to contribute to (inter)disciplinary conversations. (R) | |
| I'm quite good at talking about my research ideas in ways that bring other researchers into the conversation. | Bauer-adapted |
| I would rather follow my advisor's lead and let them do the networking. (R) | |
| I reach out to others who might be interested in a research idea. | |
| I can identify potential contributors who can move a research idea forward. | |
| If I have a research idea, I can speak up in my team and be listened to. | |
| I hesitate to be involved in projects that require me to work on a team where I am the only member in my discipline. (R) | Bauer-adapted |
| I do my best research when I am working on a team. | Bauer-adapted |
| I do my best research when I work alone. (R) | Bauer-adapted |
| I am comfortable leading a group of my academic peers. | |

**Supplemental Table 3. Confidence in graduate school survey.** In total, eighteen items were used to create this scale. Items that are reverse (R) coded are noted as well as the source of items. Cronbach's Alpha yielded a value of 0.882.

| Confidence in Graduate School Scale | Source |
|---|---|
| I am confident working on research ideas in my primary discipline. | |
| I have a primary discipline, but when I think about it, I am comfortable working across many disciplines. | |
| I learn quickly across academic subjects. | |
| I don't always feel comfortable reaching out for assistance/clarification from outside my primary discipline. (R) | Bauer-adapted |
| I can manage/engage in multiple ideas simultaneously. | |
| I can see how a research project might fit into a larger program of research. | |
| I can break a research idea down into manageable steps. | |
| It does not bother me if I put forward a research idea that won't work. | |
| I am afraid of having my research ideas rejected. (R) | Rosenberg-adapted |
| I am hesitant to advance research ideas that require expertise outside my primary discipline. (R) | Bauer-adapted |
| I don't feel comfortable venturing outside my primary research direction. (R) | |
| I feel uncomfortable pitching original research ideas to my advisor. (R) | |
| I have regular discussions with my advisor about how my research is beneficial to my professional development. | |
| I enjoy developing research ideas whether they lead to a scholarly product (experiment, presentation, paper) or not. | |
| If I do not have a scholarly product to show (experiment, presentation, paper), then I have failed. (R) | |
| On the whole, I am satisfied with my scholarly progress. | Rosenberg-adapted |
| I have a number of good qualities that I contribute to my research team. | Rosenberg-adapted |
| I feel that I have a lot to contribute to my research team, at least as much as others. | Rosenberg-adapted |
